# Eye movements enhance stereoscopic 3D vision

**DOI:** 10.64898/2026.10.20.760568

**Authors:** Y. Howard Li, Jie Z. Wang, Janis Intoy, Jonathan D. Victor, Michele Rucci

## Abstract

In moving organisms, sensory systems must recover spatial properties of the world from signals continually altered by their own movements. These self-generated temporal signals carry useful spatial information across various sensory modalities^1–5^, but it remains unclear whether they also contribute to fine visual computations traditionally thought to rely on the spatial structure of the retinal images. In humans, stereopsis estimates depth from differences between the images in the two eyes with extraordinary precision^6,7^. Yet fixational eye movements continually alter these differences, causing binocular disparity to fluctuate over time^8^. Here we show that stereopsis critically depends on these oculomotor-induced input changes. Using high-resolution binocular eye tracking and gaze-contingent display control, we selectively manipulated the retinal consequences of the differential (vergence) and common (version) components of binocular eye movements. Suppressing eye-movement-induced disparity modulations severely impairs stereopsis, even when retinal image motion in each eye is preserved. Across viewing conditions, the strength of disparity modulations closely predicts performance, regardless of whether they are generated by vergence or version. These findings establish stereopsis as a spatiotemporal process and reveal disparity dynamics induced by eye movements as a critical resource for fine depth perception.

---

Textbook descriptions of stereopsis rarely consider that binocular disparity—the difference between corresponding features in the images on the two retinas—continually changes over time^6,9,10^. Binocular dispaity flucutates because the human eyes are always in motion, even during fixation (Fig. 1*a − a*), when rapid, small gaze shifts (microsaccades) alternate with a slower, meandering motion (ocular drift)^11–14^. Importantly, the intersaccadic fixational movements of the two eyes are largely independent^8,15–18^ (Fig. 1*d*), giving rise to both common (version-induced) and opposite (vergence-induced) motion of the retinal images (Fig. 1*e*), and consequent fluctuations of binocular disparity over time. Under normal viewing, disparity fluctuations can reach tens of arcminutes, far exceeding stereoacuity thresholds^7^ (Fig. 1*f* ).

**Figure 1:**
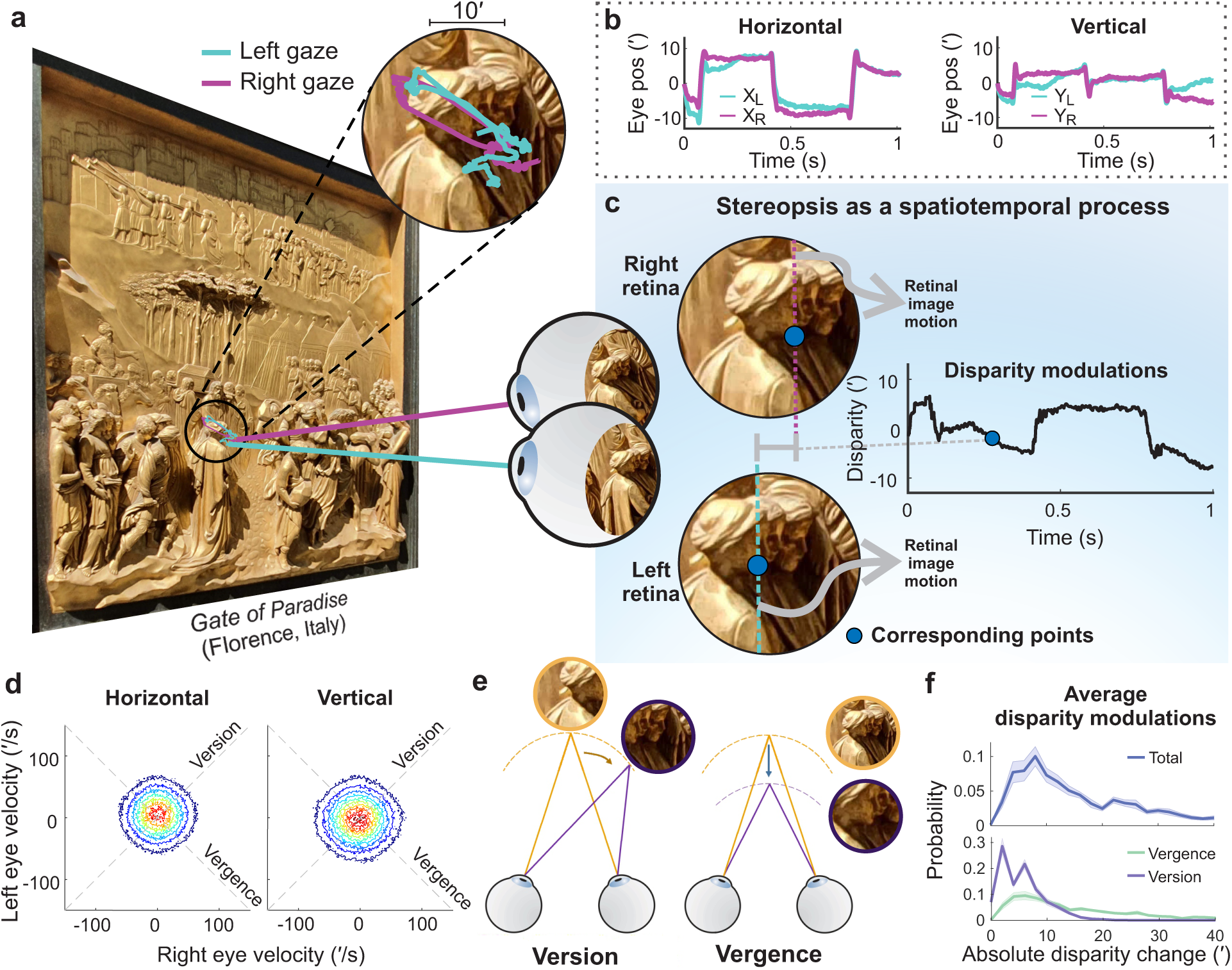
Stereopsis as a spatiotemporal process. (**a-c**) Stereopsis is traditionally regarded as a spatial process based on differences between the two retinal images (binocular disparity). However, fixational eye movements (insert in *a*), including microsaccades and ocular drift, continually shift the retinal images (*b*), causing the disparity of corresponding points to fluctuate over time (*c*). (**d**) Horizontal and vertical drift velocities are only weakly correlated between the two eyes when immobilizing the head. Data are from a representative observer. (**e**) Version and vergence generate temporal changes in disparity signals through distinct mechanisms. Version moves gaze along the zero-disparity surface (dashed arc), producing disparity changes according to variations in scene depth, whereas vergence shifts the fixation plane in distance, producing disparity changes proportional to changes in the angle between the two eyes. (**f**) Distribution of disparity changes during natural head-free fixation. For each fixation of the 3-D scene shown in (*a*), absolute deviations from the median disparity were computed and then averaged across observers (*N* =13). Total disparity change reflects the sum of disparity change from version and vergence.

These self-generated temporal changes are commonly regarded as disturbances that the visual system must cope with^19–21^. Classical studies showed that binocular fusion can persist despite substantial changes in disparity^22,23^, and that depth percepts can be elicited by very brief stimulus presentations^24,25^ or by their stereoscopic afterimages ^26,27^, suggesting that on-going eye movements are not required for stereopsis. Furthermore, it is well established that both neuronal and perceptual sensitivity are greater for static or slowly changing depth stimuli than for rapidly changing ones^28,29^. These observations support the view that stereopsis is highly robust to temporal fluctuations introduced by eye movements and primarily driven by relatively stable spatial disparities.

Yet there are reasons to question this view. Several studies have shown that stereopsis can operate at remarkably fine temporal scales^30^, with interocular delays shorter than 1 ms sufficient to elicit a percept of motion in depth^31–33^. Furthermore, fixational eye movements are increasingly recognized as playing an important role in visual processing^21,34,35^, transforming spatial patterns into temporal luminance modulations that enhance fine spatial vision^34,36,37^. Together, these findings suggest a long-standing yet under-appreciated alternative possibility^38^: rather than merely discounting the disparity fluctuations generated by eye movements, the visual system may exploit their temporal structure to support fine depth perception.

Classical studies of stereopsis that used brief flashed presentations under retinal stabilization^22,26,39,40^ did not directly test this hypothesis. The persistence of stereopsis in these studies does not rule out a possible role for self-generated temporal fluctuations, because brief flashes, which are powerful but highly unnatural stimuli for the visual system, necessarily introduce substantial temporal transients. One retinal-stabilization experiment directly compared stereoacuity with and without fixational eye movements and found similar improvements in stereoacuity with increasing exposure duration in both viewing conditions^41^. However, the stabilization method used in this pioneering experiment eliminated oculomotor-induced retinal image motion altogether, suppressing both its binocular and monocular consequences. Whether the binocular disparity modulations produced by fixational eye movements contribute to stereoscopic depth perception thus remained unresolved. Recent technological advances now allow direct examination of the functional role of these oculomotor-induced disparity dynamics in stereopsis. Here we combined high-resolution binocular eye tracking with precise gaze-contingent display control using a custom apparatus^42^. This novel setup enables high-quality retinal stabilization, and crucially, provides the flexibility to manipulate the binocular effects of self-generated retinal image motion while preserving its monocular consequences. Using this approach, we tested whether temporal disparity signals generated by fixational eye movements contribute to stereoscopic depth perception.

## Stereopsis as a spatiotemporal process

(**a-c**) Stereopsis is traditionally regarded as a spatial process based on differences between the two retinal images (binocular disparity). However, fixational eye movements (insert in *a*), including microsaccades and ocular drift, continually shift the retinal images (*b*), causing the disparity of corresponding points to fluctuate over time (*c*). (**d**) Horizontal and vertical drift velocities are only weakly correlated between the two eyes when immobilizing the head. Data are from a representative observer. (**e**) Version and vergence generate temporal changes in disparity signals through distinct mechanisms. Version moves gaze along the zero-disparity surface (dashed arc), producing disparity changes according to variations in scene depth, whereas vergence shifts the fixation plane in distance, producing disparity changes proportional to changes in the angle between the two eyes. (**f**) Distribution of disparity changes during natural head-free fixation. For each fixation of the 3-D scene shown in (*a*), absolute deviations from the median disparity were computed and then averaged across observers (*N* =13). Total disparity change reflects the sum of disparity change from version and vergence.

## Retinal motion enhances stereoscopic performance

To investigate whether the visual signals generated by fixational eye movements contribute to stereoscopic depth perception, we tested human observers’ ability to discriminate depth patterns embedded in random-dot stereograms^43^, in which the corrugation depth pattern emerged only through binocular disparity (Fig. 2*a*). Stimuli were viewed binocularly either with normal retinal motion (*normal* condition, Fig. 2*b*) or under retinal stabilization which eliminated the retinal image motion produced by fixational eye movements (*stabilized* condition, Fig. 2*c*). Retinal stabilization was achieved using a custom apparatus that integrates high-resolution binocular eye-tracking with real-time gaze-contingent display control^42,44^. On each trial, observers reported whether a sinusoidal corrugation was titled to the left or right (Fig. 2*d*).

**Figure 2:**
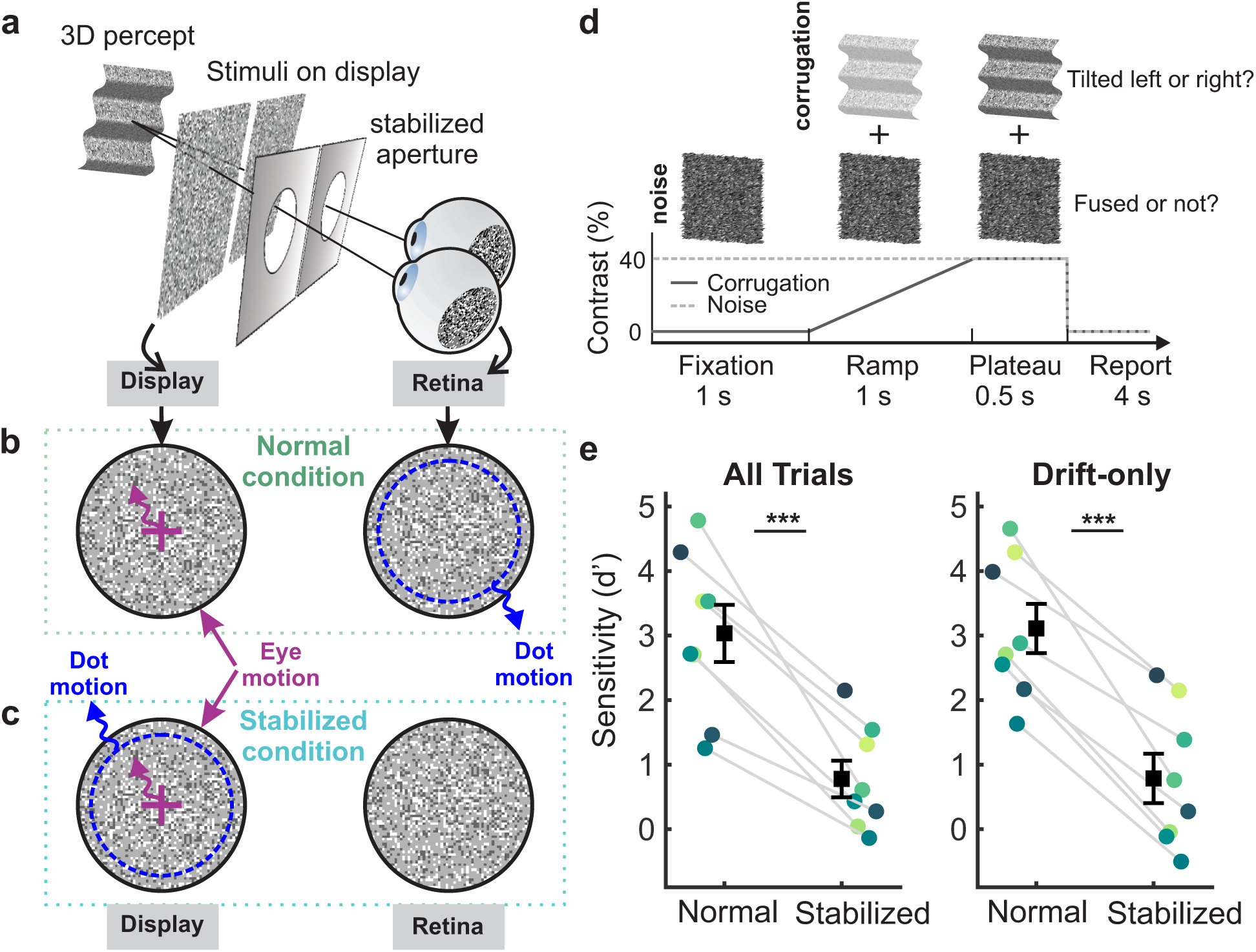
Stereoscopic depth perception depends on retinal image motion. (**a-d**) Experimental paradigm. (*a*) Observers (*N* = 8) reported the orientation (*±*10*^◦^*) of a sinu-soidal depth corrugation presented behind a gaze-stabilized aperture (radius = 5*^◦^*). (*b*) In the *Normal* condition, retinal image motion of the stimuli arose from fixational eye movements. (*c*) In the *Stabilized* condition, a gaze-contingent display updated the motion of the dots (blue arrows) in real time according to eye movements (magenta arrows), effectively immobilizing the retinal image in each eye and consequently eliminating the disparity modulations. (*d*) The corrugation was presented with gradually increasing contrast, together with a noise patch, for 1.5 s before observers reported its orientation. (**e**) Mean stereoscopic performance (*N* = 8). Stabilization reduced sensitivity by *∼*80% relative to normal viewing when all trials were included (left: *^∗∗∗^p <* 0.001, paired *t* -test). A similar reduction was observed when analyses were restricted to drift-only trials (right: *^∗∗∗^p <* 0.001, paired *t* -test). Error bars indicate *±* one SEM. Circles represent individual observers.

Strikingly, retinal stabilization substantially impaired stereoscopic performance (Fig. 2*e*). Observers readily discriminated the corrugations during normal viewing (*d^′^*=3.03 *±* 0.44, *N* = 8), whereas the discriminability dropped to 0.78 *±* 0.28 under retinal stabilization (*p <* 0.0001, paired *t* -test). This reduction was consistent across observers, with all participants individually showing statistically significant lower *d^′^* sensitivity in the stabilized condition (*p <* 0.001, *z* -test). Moreover, this impairment did not depend on microsaccades: it was present both when all trials were included (Fig. 2*e*, left) and when the analysis was restricted to trials containing only ocular drift (Fig. 2*e*, right). Thus, the presence of retinal motion normally generated by fixational eye movements can enhance stereoscopic performance.

## Stereoscopic depth perception depends on retinal image motion

(**a-d**) Experimental paradigm. (*a*) Observers (*N* = 8) reported the orientation (*±*10*^◦^*) of a sinusoidal depth corrugation presented behind a gaze-stabilized aperture (radius = 5*^◦^*). (*b*) In the *Normal* condition, retinal image motion of the stimuli arose from fixational eye movements. (*c*) In the *Stabilized* condition, a gaze-contingent display updated the motion of the dots (blue arrows) in real time according to eye movements (magenta arrows), effectively immobilizing the retinal image in each eye and consequently eliminating the disparity modulations. (*d*) The corrugation was presented with gradually increasing contrast, together with a noise patch, for 1.5 s before observers reported its orientation. (**e**) Mean stereoscopic performance (*N* = 8). Stabilization reduced sensitivity by *∼*80% relative to normal viewing when all trials were included (left: *^∗∗∗^p <* 0.001, paired *t* -test). A similar reduction was observed when analyses were restricted to drift-only trials (right: *^∗∗∗^p <* 0.001, paired *t* -test). Error bars indicate *±* one SEM. Circles represent individual observers.

## Modulations of disparity, rather than of luminance, enhance stereoscopic performance

Having established that eliminating retinal image motion impairs stereopsis, we next asked whether this effect resulted from the loss of binocular disparity modulations or from the loss of monocular luminance modulations in each eye, which are known to support visual acuity^35,37^ and discriminability^34,45,46^ under normal viewing. To distinguish these possibilities, we exploited the different effects of fixational vergence and version on luminance and disparity signals. During normal fixation of the corrugation stimulus, the two components generated retinal image motion of comparable strength in each eye and therefore similar luminance modulations. Critically, however, their effects on disparity differed markedly: vergence changes binocular disparity independently of scene content, producing substantially greater disparity modulations than version, whose effects were negligible given the small depth variations (1*^′^*) in the corrugation pattern (Fig. 3a − b).

**Figure 3:**
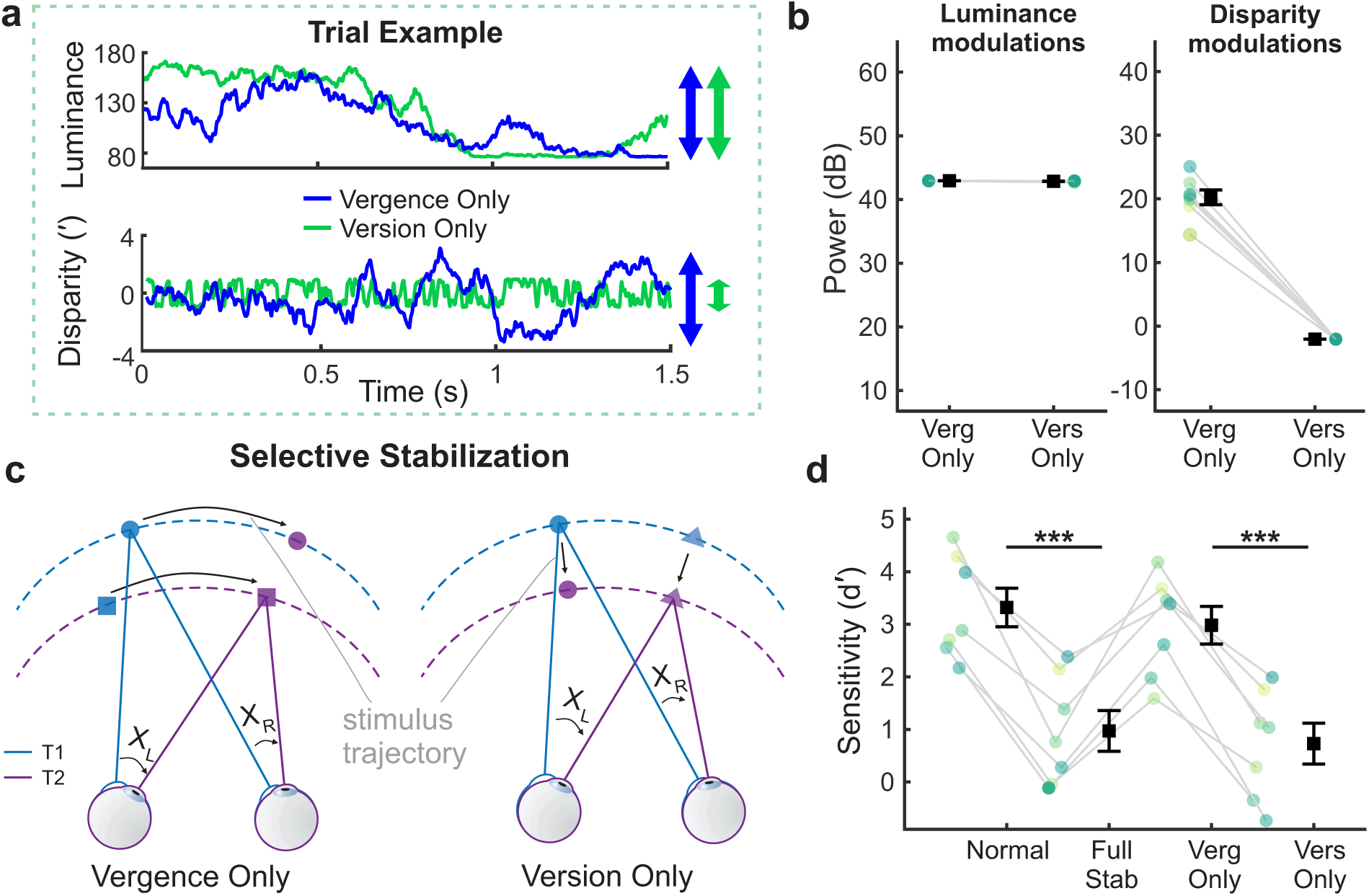
Importance of disparity modulations. (**a-b**) With the RDS corrugations of Fig. 2a, fixational version and vergence eye movements generate similar luminance modulations (top) but version consistently generated substantially weaker disparity power than vergence (bottom). (*a*) Trial example and (*b*) mean temporal power of luminance and disparity modulations across observers (*N* = 8) during normal fixation. (**c**) Schematic of selective stabilization. From *T* 1 to *T* 2, the eyes undergo a common rightward motion together with convergence. Dot positions are updated on the displays according to one component of binocular eye motion via gaze-contingent control. In the Vergence-Only condition (left), dots on different horopters (square and circle) move along their respective horopters with the common eye motion. In the Version-Only condition (right), dots on the same horopter (circle and triangle) shift to a nearer horopter following the convergence of the two eyes. (**d**) Mean stereoscopic performance (*N* = 7) under the selective stabilization conditions. Sensitivity declined significantly in the Version-Only condition (*^∗∗∗^p* = 0.006, paired *t* -test), but remained comparable to normal viewing in the Vergence-Only condition. Error bars indicate *±* one SEM. Circles represent individual observers. For comparison, results from *Normal* and *Full Stab* from Fig. 2e are here replotted for participants in both experiments. Analyses are restricted to drift-only trials in all panels.

In a further experiment, we therefore selectively preserved the retinal image motion from either version or vergence via precise gaze-contingent control: stimulus motion was yoked to one component of binocular eye motion, canceling its retinal consequences and leaving those of the other. Thus, in Vergence-Only condition, the stimulus followed the common motion of the eyes which shifts binocular gaze approximately along the same horopter, whereas in the Version-Only condition it followed their differential motion which changes vergence angle and shifts the horopter in depth (Fig. 3*c*). As expected, the two conditions retained comparable retinal image motion (*p >* 0.4, Tukey’s HSD post hoc test; Fig. S1) and associated luminance modulations.

Despite their similar retinal motion, the two conditions produced markedly different stereoscopic performance (Fig. 3*d*). *d^′^* sensitivity recovered to approximately normal levels in the Vergence-Only condition, when disparity modulations were present, but remained strongly impaired in the Version-Only condition, when disparity modulations were absent. These findings indicate that the impaired performance under retinal stabilization cannot be explained simply by the loss of luminance modulations in the individual eyes. Instead, stereoscopic performance depends critically on the temporal disparity modulations generated by binocular fixational eye movements.

## Importance of disparity modulations

(**a-b**) With the RDS corrugations of Fig. 2*a*, fixational version and vergence eye movements generate similar luminance modulations (top) but version consistently generated substantially weaker disparity power than vergence (bottom). Trial example (*a*) and mean temporal power (*b*) of luminance and disparity modulations across observers (*N* = 8) during normal fixation. (**c**) Schematic of selective stabilization. From *T* 1 to *T* 2, the eyes undergo a common rightward motion together with convergence. Dot positions are updated on the displays according to one component of binocular eye motion via gaze-contingent control. In the Vergence-Only condition (left), dots on different horopters (square and circle) move along their respective horopters with the common eye motion. In the Version-Only condition (right), dots on the same horopter (circle and triangle) shift to a nearer horopter following the convergence of the two eyes. (**d**) Mean stereoscopic performance (*N* = 7) under the selective stabilization conditions. *d^′^* sensitivity declined significantly in the

Version-Only condition (*^∗∗∗^p* = 0.006, paired *t* -test), but remained comparable to normal viewing in the Vergence-Only condition. Error bars indicate *±* one SEM. Circles represent individual observers. For comparison, results from *Normal* and *Full Stab* from Fig. 2*e* are here replotted for participants in both experiments. Analyses are restricted to drift-only trials in all panels.

## The strength of disparity modulations predicts stereoscopic performance

The finding that stereoscopic performance recovers with vergence but not version movements raises an important question: is stereoscopic performance preserved with vergence because vergence generated stronger disparity modulations, or because vergence-related retinal motion is itself uniquely exploited by the visual system? Distinguishing between these possibilities requires testing whether version can support stereopsis when it generates substantial disparity modulations.

We therefore modified the stimulus in this third experiment to systematically increase the disparity modulations from version for each observer. We replaced the corrugation stimulus with a planar surface slanted in depth (Fig. 4*a*). For this stimulus, the disparity modulations generated by version increased with the slant angle *θ*, while the luminance modulations produced by vergence and version remained comparable, as were the disparity modulations produced by vergence. For each observer, we selected a slant angle that increased the disparity gradient across the visual field, allowing version to generate disparity modulations comparable to or greater than those produced by vergence (Fig. 4*b − b*). Observers reported whether the top or bottom of the slant surface was tilted toward them.

**Figure 4:**
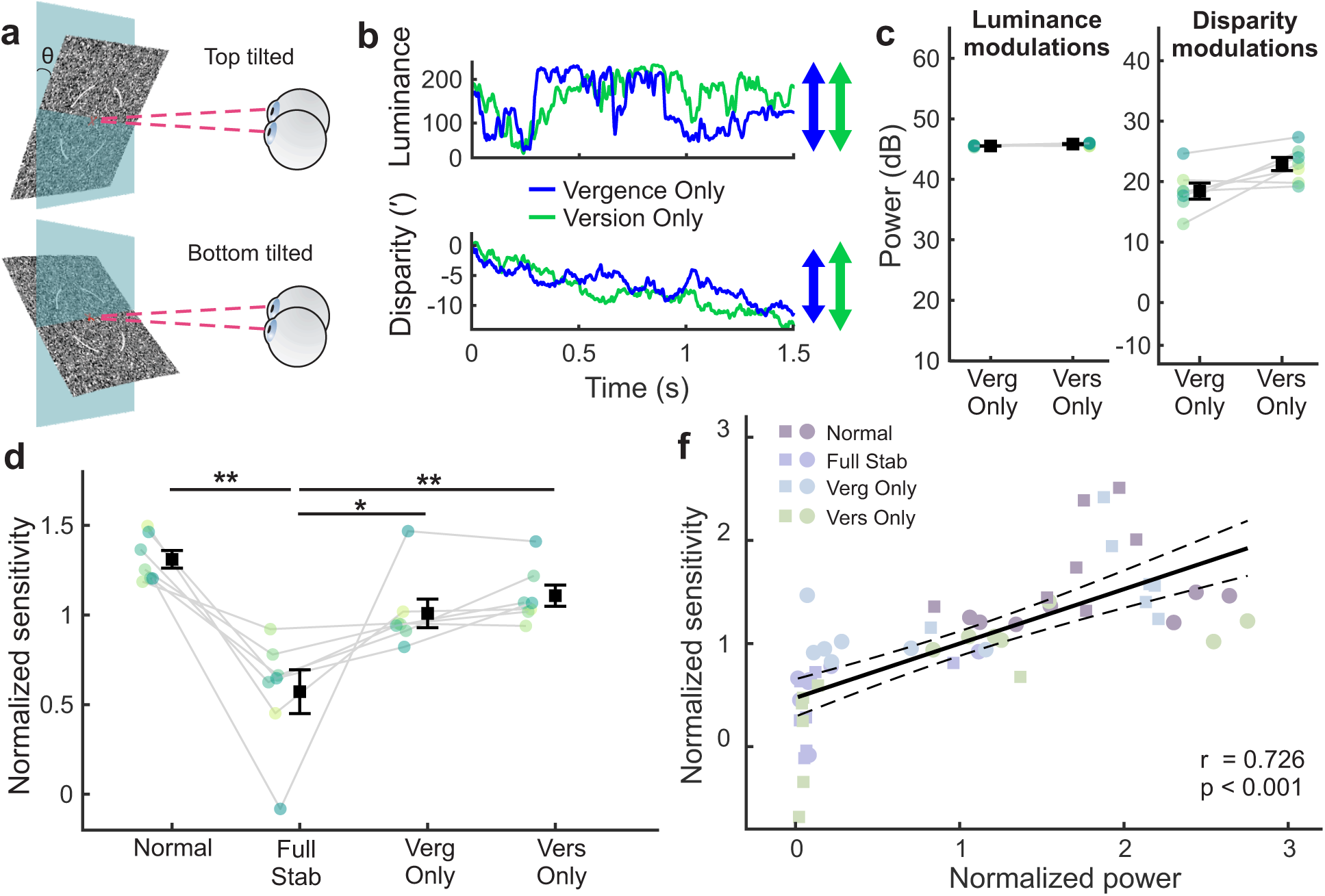
The strength of disparity modulations predicts stereoscopic performance. (**a**) Observers discriminated whether the top or bottom of a slanted surface appeared closer in depth while maintaining central fixation. (**b**) Fixational vergence and version generated comparable luminance and disparity modulations. Results were estimated from representative eye traces superimposed on the slanted surface. (**c**) Mean temporal power (*N* = 7) of luminance and disparity modulations during normal fixation of the slanted surface. The slant angle (*θ*) was individually adjusted for each observer so that version-induced disparity modulations were comparable to or greater than vergence-induced disparity modulations. Luminance modulations remained comparable between version and vergence. (**d**) Mean stereoscopic performance (*N* = 7) under four viewing conditions. Full retinal stabilization (*^∗∗^p <* 0.001) reduced sensitivity, whereas preserving either vergence-(*^∗^p* = 0.021) or version-(*^∗∗^p* = 0.005) disparity modulations restored performance (one-way ANOVA with post hoc comparison tests). (*e*) (**f**) Across both corrugation (circle markers) and slant experiments (square markers), stereoscopic performance increased with the strength of disparity modulations (*^∗∗^p <* 0.001). Each color denotes a viewing condition. Analyses were restricted to drift-only trials in all panels.

Performance closely followed the strength of disparity modulations (Fig. 4*d*). Full stabilization yielded the lowest *d^′^* sensitivity among the four viewing conditions, whereas preserving either vergence- or version-induced disparity modulations substantially improved performance. Thus, unlike in the corrugation experiment, version alone supported robust stereoscopic performance when it generated sufficiently strong disparity modulations. Neither the Vergence-Only nor Version-Only condition, however, reached the sensitivity observed during normal viewing (*p <* 0.02). This is expected if both components contribute to the temporal disparity signals available during normal fixation, such that eliminating either component reduces their overall strength.

To further corroborate this hypothesis, we examined whether stereoscopic performance across viewing conditions could be predicted by the magnitude of these signals. As shown in Fig. 4*f* , stereoscopic performance was strongly correlated with the temporal power of disparity modulations across both experiments (Pearson r = 0.726, p *<* 0.001).

Together, these findings show that stereoscopic performance is determined primarily by the strength of the temporal disparity modulations generated by fixational eye movements, rather than by whether those modulations arise from vergence or version. The poor performance in the Version-Only condition of the corrugation experiment of Fig. 2*d* therefore reflected the weakened disparity modulations available in that condition, rather than the absence of vergence per se.

## The strength of disparity modulations predicts stereoscopic performance

(**a**) Observers discriminated whether the top or bottom of a slanted surface appeared closer in depth while maintaining central fixation. (**b**) Fixational vergence and version generated comparable luminance and disparity modulations. Results were estimated from representative eye traces superimposed on the slanted surface. (**c**) Mean temporal power (*N* = 7) of luminance and disparity modulations during normal fixation of the slanted surface. The slant angle (*θ*) was individually adjusted for each observer so that version-induced disparity modulations were comparable to or greater than vergence-induced disparity modulations. Luminance modulations remained comparable between version and vergence. (**d**) Mean stereoscopic performance (*N* = 7) under four viewing conditions. Full retinal stabilization (*^∗∗^p <* 0.001) reduced *d^′^* sensitivity, whereas preserving either vergence-(*^∗^p* = 0.021) or version-(*^∗∗^p* = 0.005) disparity modulations restored performance (one-way ANOVA with post hoc comparison tests). (*e*) (**f**) Across both corrugation (circle markers) and slant experiments (square markers), stereoscopic performance increased with the strength of disparity modulations (*^∗∗^p <* 0.001). Each color denotes a viewing condition. Analyses were restricted to drift-only trials in all panels.

### Discussion

The finding that oculomotor-induced disparity modulations enhance fine depth perception challenges the conventional characterization of stereopsis as primarily a spatial computation^41^. Stereopsis instead benefits from temporal structure that fixational eye movements impose on retinal disparities. Importantly, this benefit cannot be explained by the known monocular consequences of retinal image motion, as preserving retinal motion in each eye without strong disparity modulations did not restore stereoscopic performance.

Our results further show that the availability of relative disparity cues alone does not determine stereoscopic performance. Relative disparity has long been regarded as the most reliable cue for stereoscopic vision, whereas absolute disparity alone produces comparatively weak depth percepts^47–50^. These experiments, along with the theoretical advantage that the relative disparity signal is invariant to eye movements, have contributed to the prevailing view that fine depth perception relies predominantly on relative disparity^10^. In our experiments, however, relative disparity cues were unchanged across viewing conditions, whereas stereoscopic performance differed systematically with the temporal modulation of absolute disparity. Thus, the extraction or use of relative disparity appears to be facilitated by dynamic absolute disparity signals.

Our results demonstrate that disparity modulations generated by both vergence and version are beneficial to stereopsis. However, the present study did not examine whether these benefits depend on the contingency between disparity modulations and the eye movements that generate them. One possibility is that the visual system combines retinal disparity dynamics with extra-retinal information about binocular eye motion^51^, and uses this extraretinal information to compute depth. Under this account, consistency between oculomotor events and visual inputs could contribute to stereoscopic processing^52^. Alternatively, temporal disparity modulations may be exploited directly, irrespective of whether they arise from self-generated or externally imposed image motion, analogous to the temporal luminance modulations generated by ocular drift^34,37^. Distinguishing these possibilities will require comparing the effectiveness of self-generated disparity modulations with externally generated signals matched in their retinal dynamics.

Together, our findings extend to stereopsis a principle observed across other sensory systems, that is, self-generated temporal signals can provide spatial information useful for perception. Fixational eye movements do not merely perturb retinal disparity; they generate temporal structure that supports fine depth judgments. Thus, even visual computations traditionally regarded as primarily spatial can exploit signals generated by the observer’s own movements. Future work will need to determine how broadly these dynamic mechanisms operate in natural vision and how they interact with other sensory cues to support a coherent percept of three-dimensional space.

### Methods

## Subjects

Fourteen emmetropic observers participated in the experiments of this study (6 males; age range: 18-30 years). Eight observers, including author JI, participated in the Corrugation experiment, while the remaining seven observers, including authors HL and JI, participated in the Slant experiment. Data from one of these observers were excluded from analyses shown in Fig. 3 because the observer did not complete all viewing conditions. With the exception of the participating authors, all observers were näıve to the purpose of the study. All participants provided informed consent before participation and näıve observers received monetary compensation for their participation. The study protocol followed the ethical procedures approved by the Institutional Review Board at the University of Rochester.

## Stimulus and Apparatus

The stimuli were random-dot stereograms (RDS) composed of black and white dots presented on a uniform gray background (mean luminance 13 cd/m^2^ with 40% contrast). Dot density was set to be 90 dots/deg^2^, and each dot subtended 3*^′^*. In the two Corrugation experiments, dot disparities were arranged to form a 1 cpd oriented sine-wave corrugation in depth with a disparity amplitude of 1*^′^* (corrugation; Fig. 2*a*). The corrugation was tilted by *±*10*^◦^* from horizontal. In the Slant experiment, dot disparities varied linearly with horizontal eccentricity to create a planar surface slanted about the horizontal axis (Fig. 4*a*). The slant angle *θ* was determined by the slope of the disparity gradient and individually adjusted to yield approximately 80% performance in the normal viewing condition (mean: 52*^◦^±*5.8*^◦^*). In all experiments, an equal number of dots were used to generate disparity noise, with disparities drawn randomly from a normal distribution (SD = 0.5*^′^*). A red fixation cross (10*^′^*) surrounded by arcs (1*^◦^* radius) was displayed to aid fixation.

Stimuli were presented on a mirror stereoscope with either two synchronized LCD monitors (Corrugation: ASUS ROG Swift 258, 1920*×*1080 pixel resolution, 200Hz refresh rate) or a single LCD monitor with split displays (Slant: ASUS PG259QN, 1920*×*1080 pixel resolution, 360Hz refresh rate) in a dimly illuminated room. Mirrors positioned near the eyes directed each line of sight to its corresponding display to ensure monocular stimulation for each eye. A bite-bar and head-rest were used to minimize head movements and maintain the observer in a fixed viewing position throughout the experiment. In each trial, stimuli were presented behind circular apertures which were stabilized with the gaze motions to keep the visual stimulation restricted to the central 5*^◦^*.

The horizontal and vertical positions of each eye were measured by means of a binocular Dual Purkinje Image (DPI) eye-tracker (Fourward Technology) or a custom digital Dual Purkinje Image eye-tracker (the dDPI ^42^). Both systems are capable of resolving movements as small as 1*^′^*, and yield data samples at 1kHz. Stimuli were rendered by means of Eye-RIS, a custom system for binocular gaze-contingent display control that enables real-time synchronization between eye movement recordings and the rendering of the image on the monitors^44^.

## Experimental Procedures

Each participant completed several experimental sessions, each about 1.5 hours in duration. Sessions began with setup procedures, including calibrating the eye trackers, aligning the stereoscopic displays, and verifying stable gaze-contingent stimulus presentation. The experiment was organized into blocks of 5-10 minutes, interleaved with short rest periods to minimize fatigue.

Proper alignment of the mirror stereoscope was confirmed before each block. First, a 3*×*3 grid of dots was presented to both eyes, and the participants adjusted the mirrors until the images appeared fused. To verify that each eye was receiving input solely from its corresponding monitor, participants used a joypad to show a crosshair on one display while the other showed a uniform gray field. By alternately closing one eye and viewing the contralateral display, they confirmed that monocular stimulation was isolated to the intended eye. Then, they adjusted the horizontal and vertical position of the grid in each eye using the joypad until toggling the grid on and off between the two eyes produced no perceptible motion, ensuring precise binocular alignment.

Accurate localization of the line of sight on each display was achieved through a two-stage calibration procedure as previously reported in detail^46,53^. Briefly, participants first sequentially fixated each point of a standard 3*×*3 grid presented monocularly to each eye, providing an initial estimate of the bilinear mapping between eye tracker signals and visual angles. In the second stage, gaze-contingent stage, the mapping coefficients were refined separately for each eye: participants adjusted the estimated gaze position—shown in real time for each grid point—until it was centered on their perceived point of fixation. During this step, one eye viewed the stimulus while the other was presented with a uniform gray field. This procedure localizes the stimulus point projecting to the preferred retinal locus of fixation in each eye with approximately tenfold greater accuracy than standard eye-tracking calibrations ^46^. To correct for small drifts in the apparatus or subtle head movements, the central fixation point was recalibrated using the gaze-contingent procedure after every three trials.

Observers were instructed to maintain fixation on the central red cross in each trial and reported the stimulus configuration via a joypad in a two-alternative forced-choice (2AFC) task. Observers reported the orientation of the corrugation pattern (left versus right) in the Corrugation experiments and the tilt of the slant surface (up closer or bottom closer) in the Slant experiment. Each trial began with a display of only the noise-pattern dots. After 2 s of fixation, the dots containing disparities of the stimulus appeared, their contrast ramping from 0 to 40% over 1 s and then remaining constant for 0.5 s. This ramping of contrast minimizes visual transients unrelated to eye movements^36,54^. The disappearance of the dots signaled the start of a 4 s response window during which observers reported both the orientation and whether the stimuli remained fused. Trials where fusion appeared broken were excluded from analyses.

Each trial was randomly assigned to be one of the four viewing conditions. In the Normal condition, dots remained stationary on each display and moved normally on each retina consequent to the observers’ eye movements (Fig. 2*b*). In the Stabilized condition, dots moved on each display according to the eye motion to counteract the effects of eye movements and eliminate image motion on the retina (Fig. 2*c*). In the Vergence-Only condition, dots moved on each display according to the average motion of the eyes so that the image on each retina reflected only the difference in the eye movements (*i*.e. version stabilized; Fig. 3*c*). In the Version-Only condition, dots moved on each display according to the difference in eye motion so that the image on each retina reflected only the average of the eye movements (*i*.e. vergence stabilized; Fig. 3*c*). The fixation aids remained stationary on the monitor in all conditions.

## Data Analysis

Analog oculomotor traces (DPI) were low-passed at 500 Hz and sampled at 1 kHz with automatic detection of no-tracking and blink period. The digital DPI provided time series sampled at 1 kHz directly. The sample traces were then segmented into periods of saccades and drifts using a velocity-based algorithm as previously described^35^. Instantaneous speed and curvature of the recorded traces were estimated after low-pass filtering oculomotor data with a Savitzky– Golay filter with cut-off frequency at approximately 40 Hz. After manual inspection, trials containing sub-optimal tracking, eye blinks, or in which the subject looked away from the fixation aids (*>*1*^◦^*) during stimulus presentation were discarded from data analysis. Effects are reported as *d^′^* sensitivity.

Retinal image motion was reconstructed by combining the recorded eye trajectories with the programmed motion of the stimulus in each display. For drift intervals, motion magnitude on each retina was summarized by the diffusion constant, *D* (Fig. S1*b*), obtained by regressing the variance of retinal displacement against time: *σ*^2^(*t*) ∝ 4*Dt*. This approach also extends to binocular drift analysis, as both version and vergence drifts can be modeled as Brownian processes.

Disparity modulations from binocular eye movements were calculated with the following equivalent formulas:

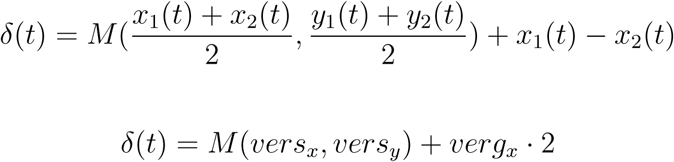

where (*x_j_*(*t*)*, y_j_*(*t*)) is the position of eye *j* at time *t* and *M* (*x, y*) denotes the disparity map of the stimulus across space. Note that disparity modulations from version and vergence combine additively, and that modulations from version depend on the stimulus *M* while modulations from vergence do not. For the Corrugation experiments, *M* (*x, y*) formed a corrugation pattern:

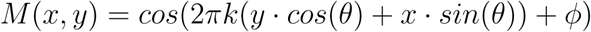

where *k* was the spatial frequency of the corrugation, *θ* was the tilt angle, and *ϕ* was the phase which was uniformly randomized between 0 and *π*. For the Slant experiment, *M* (*x, y*) formed a slanted surface:

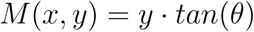

where *θ* was the slant angle along the horizontal axis.

In the example disparity modulations during natural viewing in Fig. 1*a*, stereo images of the Gates of Paradise were taken at the Baptistery of San Giovanni by author MR using a handheld camera simulating binocular viewing conditions. The disparity map (*M* (*x, y*)) of the selected panel was calculated with a Semi-Global Matching algorithm^55^, and subsequently superimposed with eye traces to produce results presented in Fig. 1*c, f* .

Spectral analyses were conducted following the approach described in previous work^56^. The spectral density of a given temporal disparity input was evaluated using Welch algorithm with 50% block overlap. This was done for each trial, and was averaged across trials for each condition and each observer. To examine the effective disparity modulation in driving visual responses, the spectral density was weighted by the temporal sensitivity of V1 neurons reported in neurophysiology literature^28^. To compare across conditions and experiments, the resulting power was normalized for each individual observer and was reported in Fig. 4*f* .

## Acknowledgments

This work was supported by grants R01 EY18363 and P30 EY001319 from the National Institutes of Health. We thank the members of the Active Perception Laboratory at the University of Rochester for helpful discussions.

## Author contributions

YHL and MR conceived the study. YHL and JS performed the study. YHL, JS and JZW analyzed the results. JZW prepared the figures and wrote the manuscript. All authors contributed to the editing of the manuscript. MR supervised the project.

## Supplementary Information

**Figure S1:**
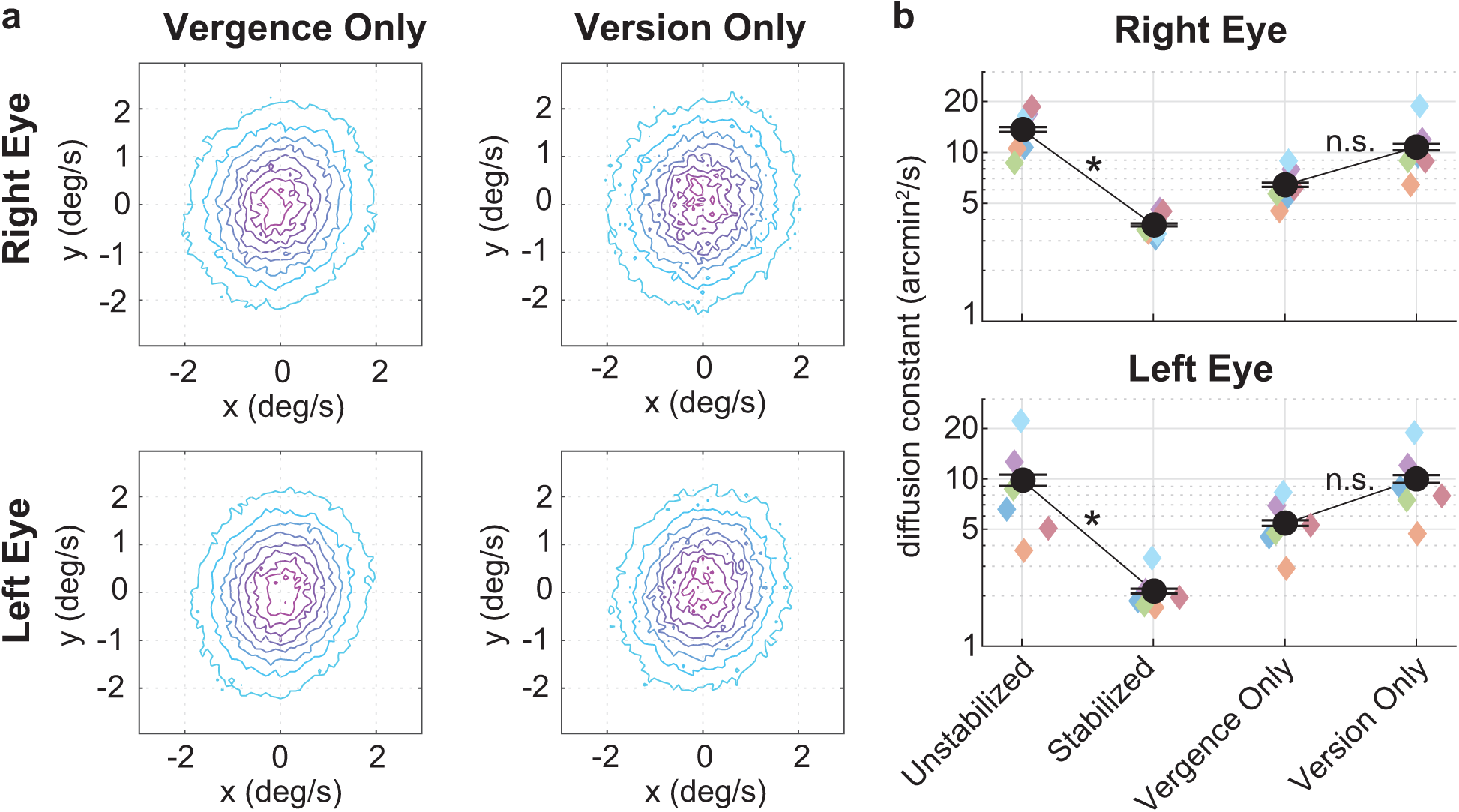
Characteristics of retinal image motion. (**a**) Distributions of retinal image velocity in each eye when only version or vergence motion is allowed. Colored contours indicate iso-probability lines and are the same across panels. Each distribution is averaged across all observers. (**b**) The diffusion constant of retinal image motion across stabilization conditions for each eye. (*^⋆^p <* 0.03, Tukey’s HSD post hoc test). Colored diamonds indicate individual subjects. Black circles and error bars are mean and standard errors across subjects.

